# Adding context to the pneumococcal core genes – a bioinformatic analysis of the intergenic pangenome of *Streptococcus pneumoniae*

**DOI:** 10.1101/2021.08.29.458057

**Authors:** Flemming Damgaard Nielsen, Jakob Møller-Jensen, Mikkel Girke Jørgensen

## Abstract

Whole genome sequencing offers great opportunities for linking genotypes to phenotypes aiding in our understanding of human disease and bacterial pathogenicity. However, these analyses often overlook non-coding intergenic regions (IGRs). By disregarding the IGRs, crucial information is lost, as genes have little biological function without expression. In this study, we present the first complete pangenome of the important human pathogen *Streptococcus pneumoniae* (pneumococcus), spanning both the genes and IGRs. We show that the pneumococcus species retains a small core genome of IGRs that are present across all isolates. Gene expression is highly dependent on these core IGRs, and often several copies of these core IGRs are found across each genome. Core genes and core IGRs show a clear linkage as 81% of core genes are associated with core IGRs. Additionally, we identify a single IGR within the core genome that is always occupied by one of two highly distinct sequences, scattered across the phylogenetic tree. Their distribution indicates that this IGR is transferred between isolates through horizontal regulatory transfer independent of the flanking genes and that each type likely serves different regulatory roles depending on their genetic context.

## Introduction

*Streptococcus pneumonia* (pneumococcus) is the leading cause of sepsis, meningitis and bacterial pneumoniae in children worldwide [1]. Widespread antibiotic resistance and the emergence of non-vaccine serotypes is making treatment increasingly difficult. These threats have led the WHO to list pneumococcus as a ‘priority’ pathogen [1-2]. This clinical relevance of pneumococcus has, in part, led to great scientific interest and the publication of several thousand sequenced genomes [3].

The availability of whole genome sequence (WGS) data has made it possible to study the entire pangenome of an organism rather than single isolates. A pangenome consists of the collected gene pool present in a group of organisms belonging to the same clade [4]. The pangenome can be divided into a core genome, which constitutes genes present in all isolates and the accessory genome as the remaining genes [4]. The pangenome of pneumococcus is considered at the extreme end of being open, that is, there is no defined limit to its pangenome as new genes are acquired continuously [5]. This openness is mainly due to new genes being acquired through horizontal gene transfer (HGT) mediated by pneumococcus’ natural competence [6-7].

Traditionally, pangenomes are limited to genes thereby excluding the non-coding intergenic regions (IGRs) [8-9]. This focus on genes alone leaves out 15% of the genomes and ignores a significant amount of crucial genomic information as IGRs contain several biologically relevant elements such as promoters, terminators, regulatory binding sites and non-coding RNAs [10-15]. To effectively link genotypes to phenotypes through pangenomics, IGRs must be taken into consideration, as genes have little biological function without expression.

Recently, IGRs have attracted more attention as potential drivers of evolution [16-18]. They persist through purifying selection, also known as negative selection, where unused or unwanted traits are removed. This persistence is true across several diverse bacterial species, in a similar fashion to that of core genes, even when major regulatory elements are excluded [16-17]. Small variations in IGRs can lead to great phenotypical impact, for instance, the inversion of a single promoter element was demonstrated to turn a commensal bacterium pathogenic [19].

IGRs may also undergo genetic recombination, a term coined *horizontal regulatory transfer* (HRT) [20-21]. HRT can occur with the flanking genes of the IGR, but in some cases, the IGRs are transferred independently of the genes they regulate [18]. As much as 32% of the core regulatory regions in *Escherichia coli* and 51% of the overall core IGRs are thought to have been acquired in this manner indicating that HRT is indeed common [18]. Another aspect of HRT is regulatory switching where one IGR is replaced with another non-homologous IGR. This leads to two or more conserved IGRs occupying the same genomic space across different isolates of the same species [18-22]. As much as 13% of the IGRs within the core genome of *E. coli* has undergone regulatory switching [18]. Thus, IGRs seemingly contribute to greater variation in the core genome than genes themselves, thereby challenging the view of the bacterial core genome as being relatively stable [18].

In this study we map the complete core genome of pneumococcus and compare the nature of genes and IGRs against each other in the pangenome. We find a clear linkage between core genes and core IGRs, but core genes are associated with different IGRs, indicating that the pneumococcal core genome is less stable than previously thought. Additionally, we identify any potential regulatory switching events within this core genome. To our knowledge we are the first to identify the complete core genome of pneumococcus, both coding and non-coding.

## Results

In this study, we map the first complete pangenome of pneumococcus, spanning both genes and IGRs. The identified intergenic core genome is provided in Appendix 1. Additionally, we screen for any regulatory switching events present within the core genome.

### Many intergenic regions are universally conserved across all pneumococcal isolates

We created a pangenome of 84 different pneumococcal isolates, spanning both genes and the non-coding IGRs. To put the nature of the pneumococcal IGR pangenome into perspective, we performed the same analysis for *S. aureus*. Both species may colonize the human upper respiratory tract, both are opportunistic pathogens and both possess open pangenomes, making them prime candidates for comparison [23]. The analysis shows that the otherwise non-coding IGRs of both species are conserved in a similar manner to genes across the pangenome, although the number of unique genes outnumber the number of unique IGRs in both species (Fig. 1).

**Fig. 1.**
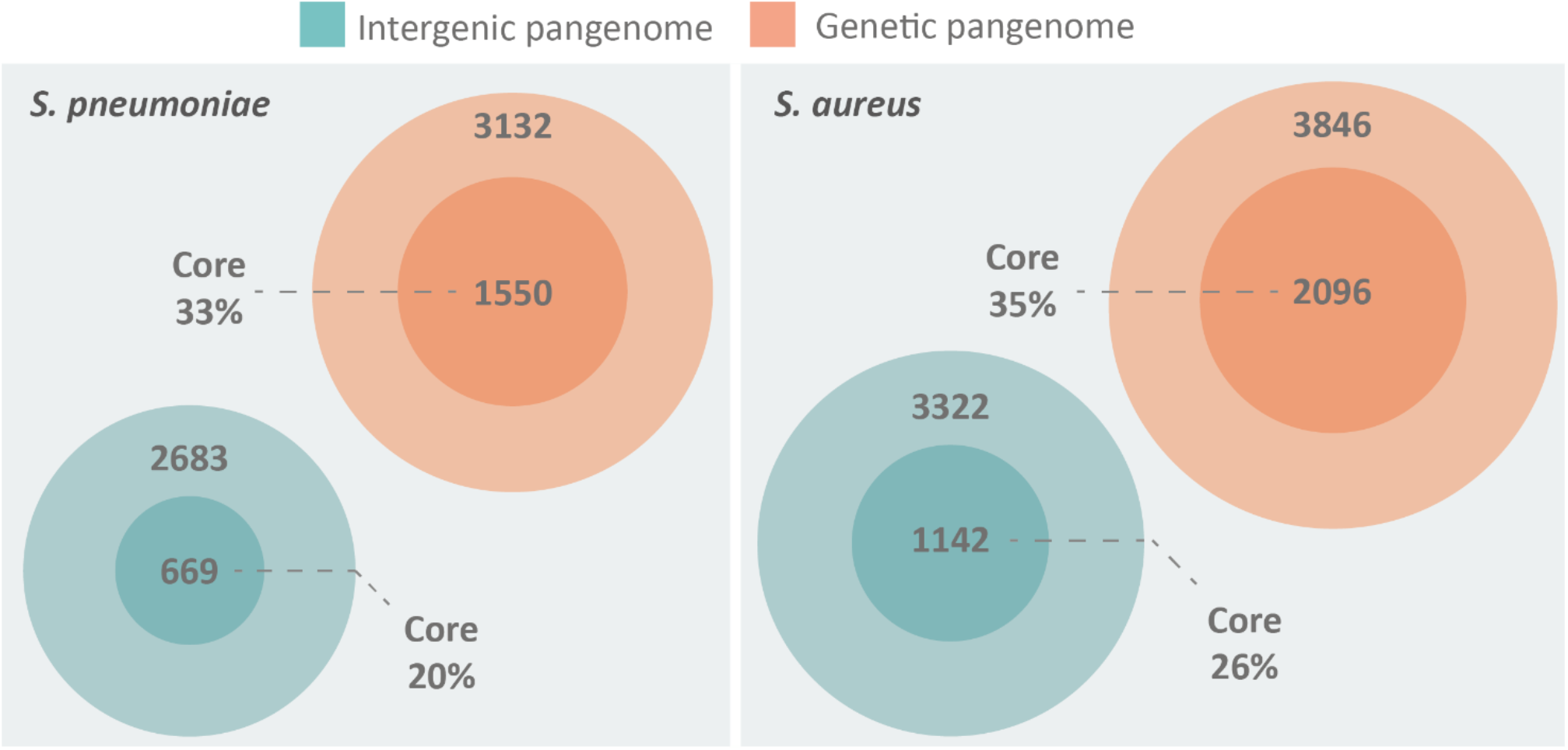
The pangenome of S. pneumoniae and S. aureus, spanning both intergenic regions (green) and genes (orange), illustrated by Venn diagrams. Both species possess a core genome of both IGRs and genes, defined as being present in >95% of isolates. The pangenomes are constructed from 84 unique genomes of each species. S. pneumoniae has a core genome of 1550 genes and 669 IGRs, while an accessory genome of 3132 genes and 2683 IGRs. S. aureus has a core genome of 2096 genes and 1142 IGRs, while an accessory genome of 3846 genes and 3322 IGRs.

While the proportion of core genes roughly scales relative to the size of the genome (pneumococcus 2.1Mbp / *S. aureus* 2.8Mbp) the proportion of core IGRs relative to genome size is lower in pneumococcus (Fig. 1). However, pneumococcus seemingly compensates for the lower number of unique IGRs by having multiple copies of several core IGRs in each genome. On average, each core IGR is present 1.23 times in each pneumococcal genome compared to 1.03 times in *S. aureus*. Each core gene is present 1.08 times in each pneumococcal genome and 1.02 times in S. aureus, this indicates that the high copy number of pneumococcal core IGRs is quite unusual.

### Core genes and IGRs constitute the majority of each genome

The average pneumococcal genome has 79% of its genes as core genes and 66% of its IGRs as core IGRs. The average *S. aureus* genome is comparatively close to that observed in pneumococcus, here core genes constitute 79% and core IGRs 68% of each genome (Table 1).

**Table 1.** The number of genes and IGRs in the core and accessory genome in selected genomes and across the collected pangenome, as well as the percentage of genes and IGRs that are core.

| Species/isolate | Core<br>genes | Core<br>IGRs | Accessory<br>genes | Accessory<br>IGRs | Percentage core<br>genes pr. genome | Percentage core<br>IGRs pr. genome |
| --- | --- | --- | --- | --- | --- | --- |
| *S. pneumoniae*<br>(Species average) | 1670 | 817 | 448 | 425 | 78.93% | 65.83% |
| *S. pneumoniae*<br>D39 | 1672 | 810 | 352 | 388 | 82.61% | 67.61% |
| *S. pneumoniae*<br>R6 | 1674 | 809 | 345 | 388 | 82.91% | 67.59% |
| *S. pneumoniae*<br>Tigr4 | 1703 | 820 | 447 | 447 | 79.21% | 64.72% |
| *S. aureus*<br>(Species average) | 2131 | 1170 | 570 | 544 | 78.98% | 68.28% |

Despite pneumococcus having fewer unique core IGRs relative to genome size than *S. aureus*, as stated earlier, their copy number is higher in each genome, thus the percentage of core IGRs pr genome is roughly equivalent in the two species (Table 1). The higher copy number of core IGRs in pneumococcus is also illustrated by the fact that 669 unique core IGRs exist in the pneumococcal core genome (Fig. 1) but on average each genome has 817 core IGRs (Table 1).

### IGRs are more likely to be unique to a few isolates than genes

The number of unique IGRs in the pneumococcal pangenome increases with the number of isolates analyzed in a similar manner to the number of unique genes (Fig. 2a). Overall, fewer unique IGRs are present in the pangenome than genes, part of this is due to the exclusion of intraoperonic IGRs of >30bp n length [22].

**Fig. 2.**
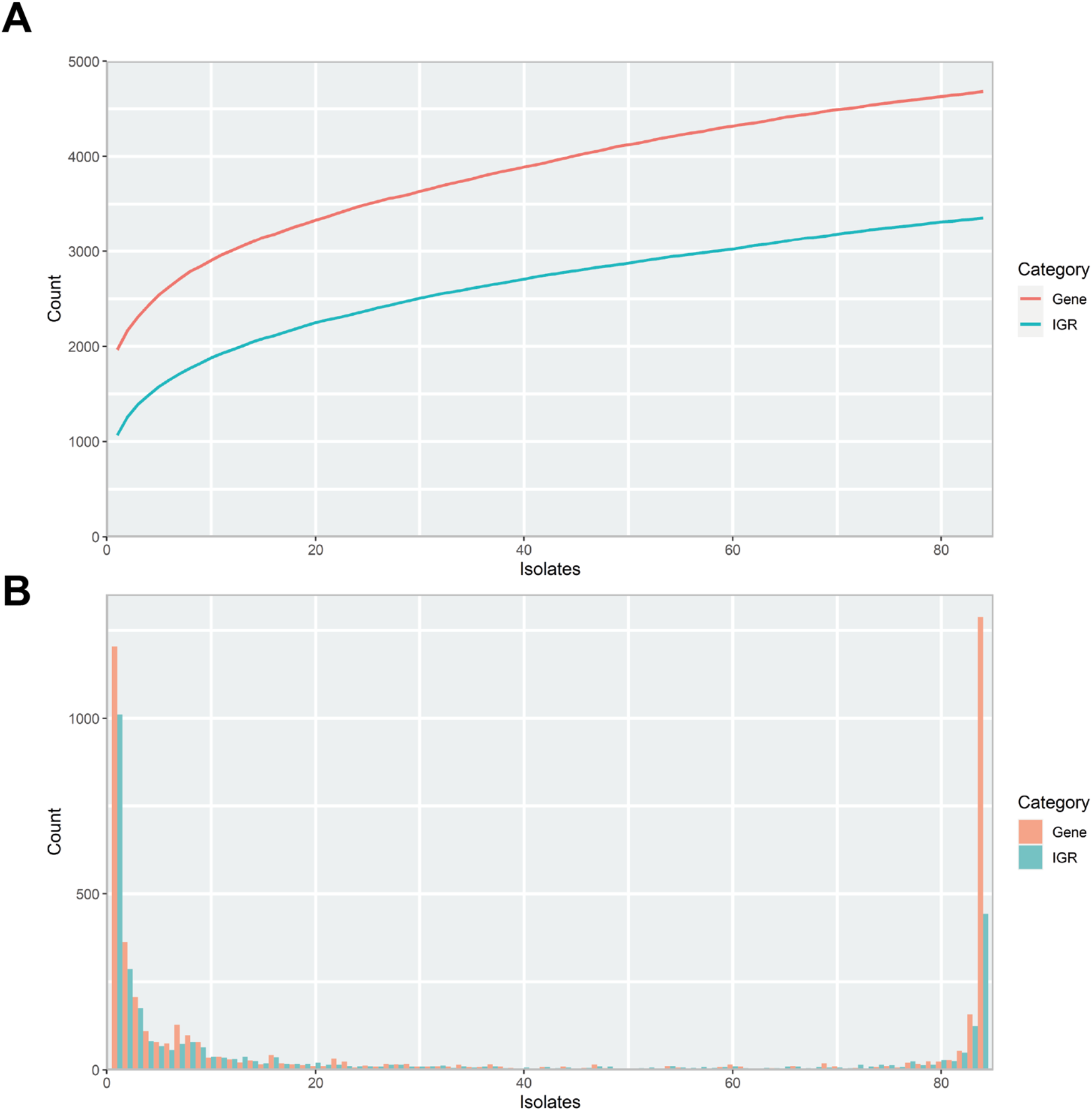
Properties of the pneumococcal pangenome and its intergenic regions (IGRs) A) Number of unique intergenic regions (green) and genes (orange) as a function of the number of isolates included in the pangenome. B) Distribution of unique IGRs (green) and genes (orange) across the streptococcal pangenome, illustrated with a frequency histogram (number of IGRs/genes present in the given number of isolates). Most IGRs and genes are part of the core genome or confined to a small fraction of the isolates.

Most IGRs are either present in almost all pneumococcus isolates or unique to only a few, that is, they are either very common or very rare (Fig. 2b). Pneumococcus genes show a similar distribution across the pangenome, though a larger proportion of IGRs are confined to only a few isolates than genes.

Pneumococcus retains more unique genes than IGRs within its pangenome (Fig. 2a), and most unique IGRs are only found in single isolates, making them rare (Fig. 2b). This scarcity of unique IGRs could indicate that IGRs experience a higher evolutionary selection threshold than genes, thereby lowering the likelihood of a newly acquired IGR of spreading to more isolates through HRT.

### Double regulatory regions are more common in the core genome

IGRs can be categorized according to the orientation of their flanking genes. IGRs that are downstream of two convergently transcribed genes are considered non regulatory (NR), IGRs that are upstream one gene and downstream another gene are considered single regulatory (SR) and IGRs that are between two divergently transcribed genes are considered double regulatory (DR) (Fig. 3).

**Fig. 3.**
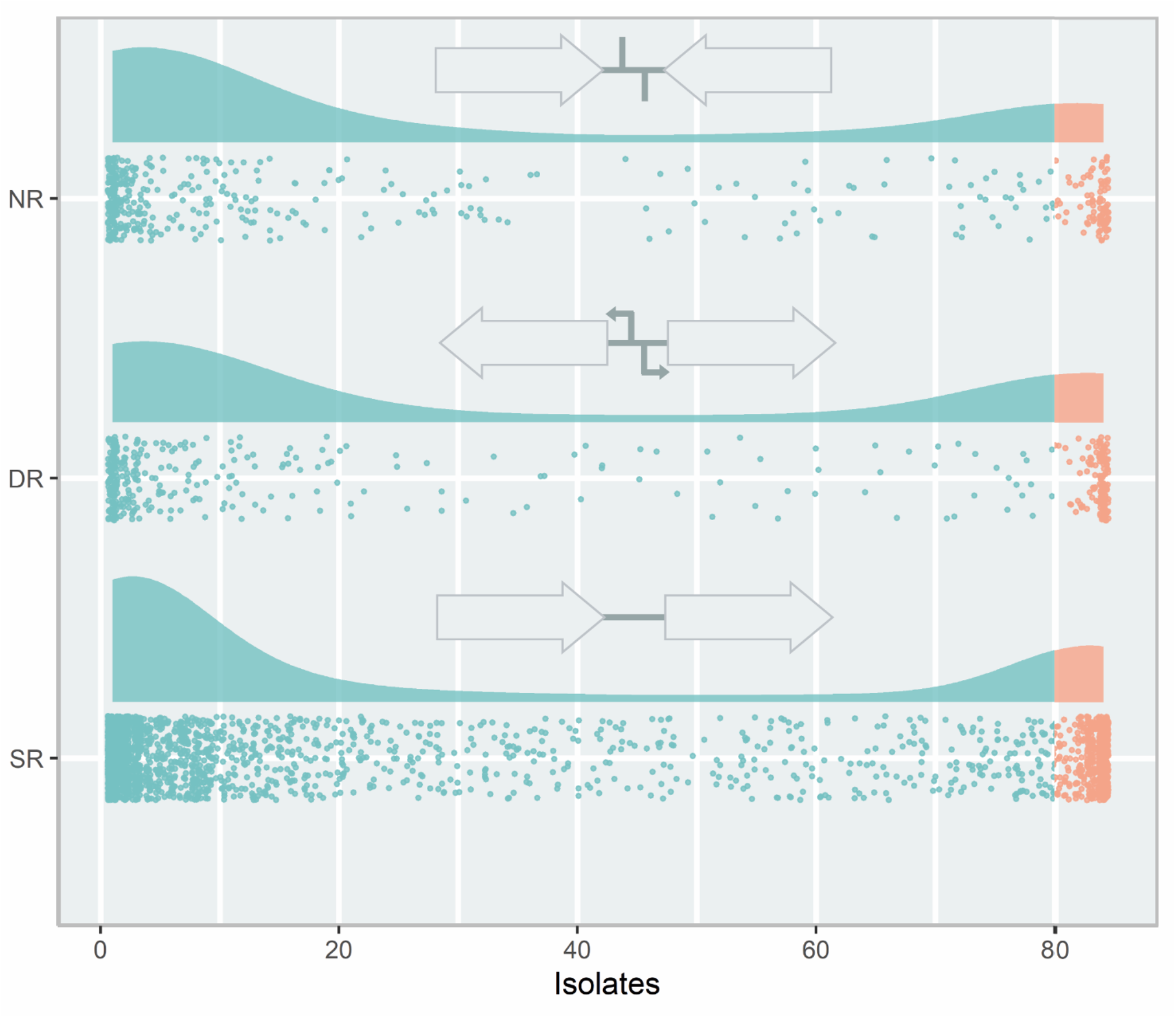
The types of intergenic regions (IGRs) and their distribution in the pneumococcal pangenome, illustrated with a raincloud plot. Each point is a unique IGR of that type plotted against the number of isolates in the pangenome it is present in. Cloud areas are scaled relative to the size of each dataset. Core IGRs are present in >95% strains (orange) and accessory IGRs are present in <95% of isolates (green). The IGRs are categorized according to the orientation of their flanking genes. If the flanking genes are pointing in the same direction the IGR is categorized as single regulatory (SR), if they face towards the IGR, it is categorized as non-regulatory (NR) and if they face away from the IGR, it is categorized as double-regulatory (DR).

Looking at the distribution of the IGR types across the pneumococcal pangenome, NR and DR regions are rare compared to SR IGRs (Fig. 3). DR regions also constitute a greater relative proportion of the core IGRs than seen in the accessory genome.

### Core IGRs are linked to core genes

Next, we analyzed the degree of linkage between core IGRs and core genes, that is, how often a core IGR is directly upstream a core gene. IGRs and their flanking genes were identified and any IGRs directly upstream the start codon of a gene was selected. The status of the IGR/gene pairs as accessory or core genome was then assessed and the ratio of each combination calculated. On average 81% of core genes in *S. pneumoniae* are associated with a core IGR, whereas only 74% of accessory genes are linked to accessory IGRs (Table 2). For comparison, the linkage of core genes to core IGRs is greater in *S. aureus* at 86%, and accessory IGRs are flanking accessory genes 82% of the time.

**Table 2.**
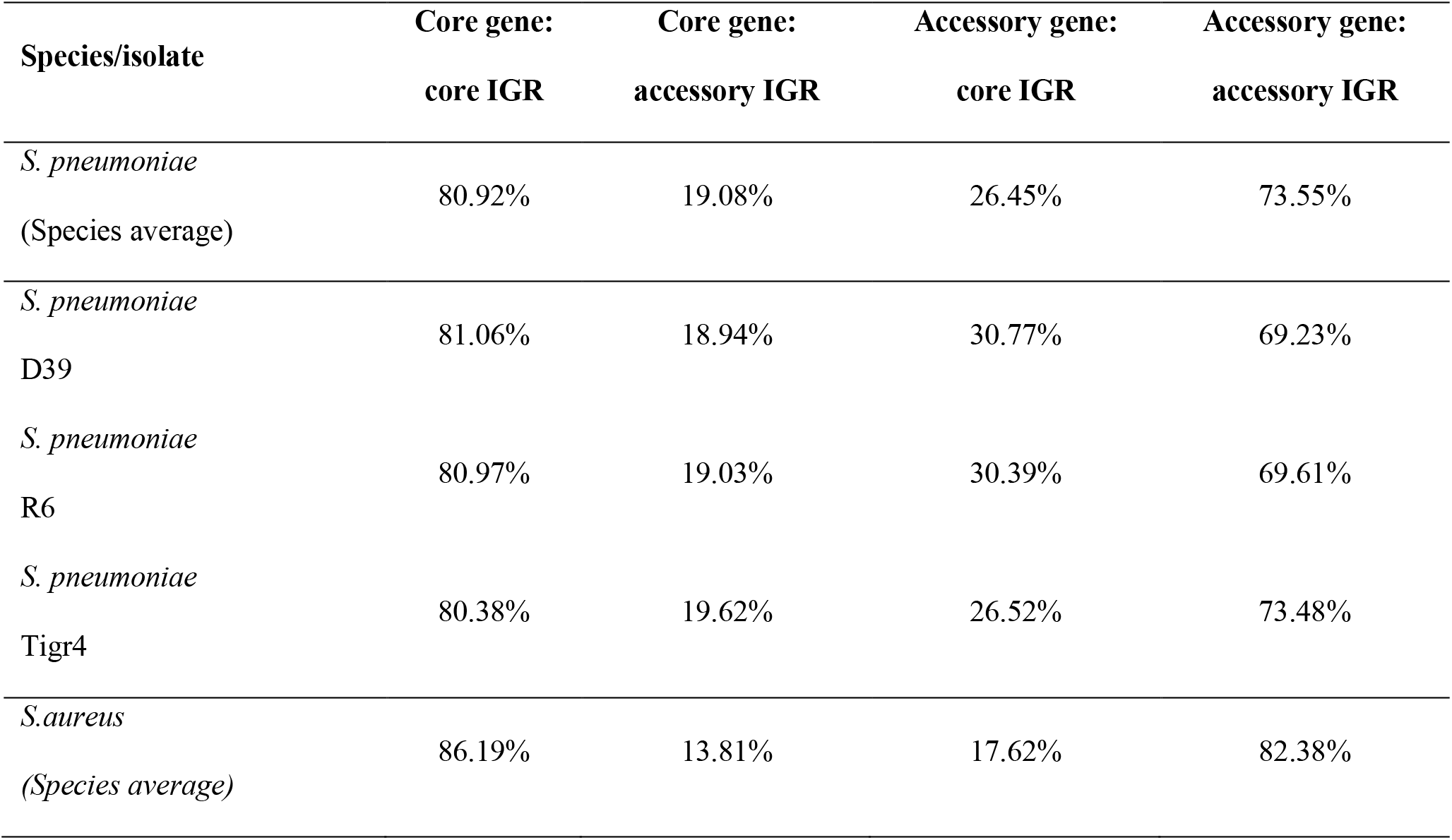
Genes and their upstream IGR was analyzed for their distribution in the pangenome. Listed are the percentage core and/or genes with a core and/or accessory IGR immediately upstream.

### A single core IGR shows sign of regulatory switching

Next, we examined the IGR candidates for regulatory switching. Regulatory switching describes when one IGR is replaced by a different non-homologue IGR. The origin of these switched IGRs is not inferred in this analysis, thus they can both originate from within the isolate itself or even from a separate species. For this analysis, only switches where the IGRs share no significant sequence homology with a BLASTN were included.

We detected three switches within the pneumococcus pangenome and only one of these is flanked by core genes. We designated the core switched IGR as csIGR (Table 3). While the two versions of the csIGR are highly conserved on their own, with both having a nucleotide identity of >99% amongst themselves, aligning the two versions with each other results in an insignificant nucleotide identity of 57%. These results were manually confirmed with a tblastn and confirmed that all pneumococcal isolates always have one of these two csIGRs but only in a single copy and always between the same flanking genes.

**Table 3.** The two versions of the csIGR in-between the single copy core genes. csIGR1 is present in 32 of the isolates analyzed and is highly conserved across the genomes with an average nucleotide identity of 99.49%. csIGR2 is present in 52 of the strains analyzed and is likewise highly conserved with an average nucleotide identity of 99.28%. Both IGRs have roughly the same length in base pairs.

|  | Length | SNPs | Nuc_identity | Length_identity | No.isolates |
| --- | --- | --- | --- | --- | --- |
| csIGR1 | 215 | 1 | 99.49% | 99.53% | 32 |
| csIGR2 | 214 | 1 | 99.28% | 99.05% | 52 |

The flanking genes were both single copy core genes and were identified in the common lab strains *S. pneumoniae* D39 and TIGR4 (Fig. 4). Interestingly, these two strains have distinct csIGR types, with D39 having csIGR1 and TIGR4 having csIGR2 (Fig. 4). Little is known about the flanking genes, other than their status as single copy core genes found in this study. SPD_1559/SP_1749 is considered essential in pneumococcus and is a homologue to *ygeH*, a gene involved in biogenesis of the 30S ribosome subunit [24-25]. The sequence of both csIGR types is provided in Appendix 2. Interestingly, neither of the csIGR types were confined to a specific phylogenetic cluster of pneumococci (Fig. 5). The fact that the csIGR types are spread across the phylogenetic tree indicates that their distribution is due to HRT. We performed a pangenome wide association study to see if any genes within the accessory genome were significantly co-occurring with the csIGR alleles across the pangenome. However, no genes were exclusively associated with neither of the csIGR types.

**Fig. 4.**
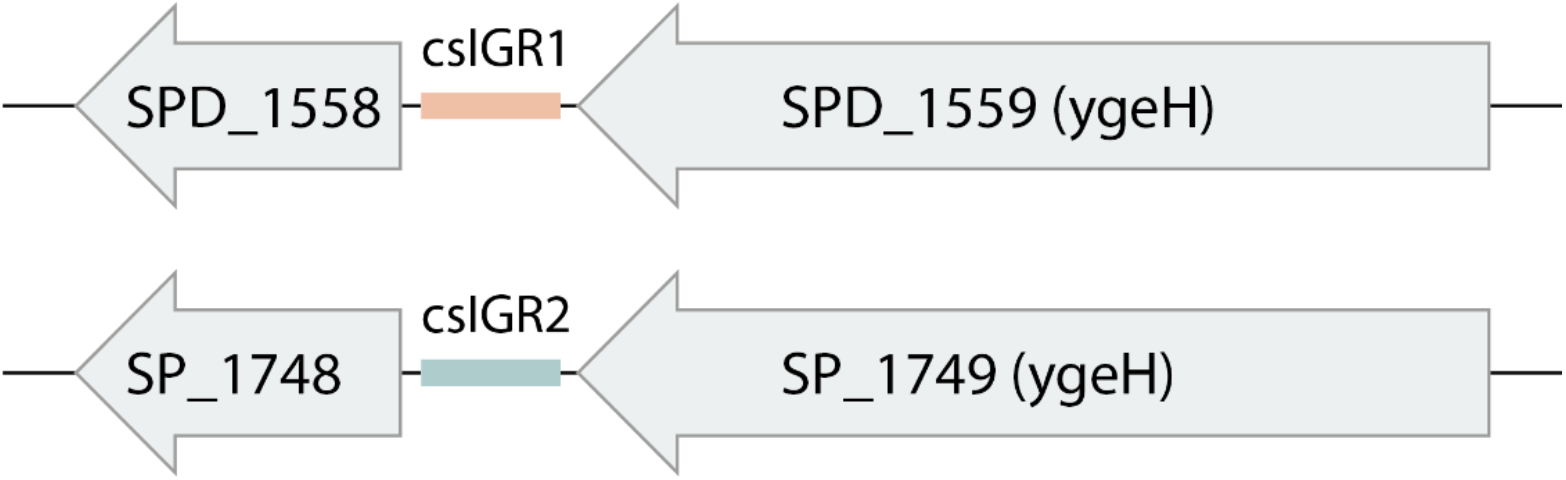
The core switched intergenic region (csIGR) in S. pneumoniae D39 and Tigr4. Each type of csIGR is represented in these strains, with D39 having csIGR1 (orange) and TIGR4 having csIGR2 (green). In D39, csIGR1 is flanked by SPD_1558 and SPD_1559. In TIGR4, csIGR2 is flanked by the genes SP_1748 and SP_1749.

**Fig. 5.**
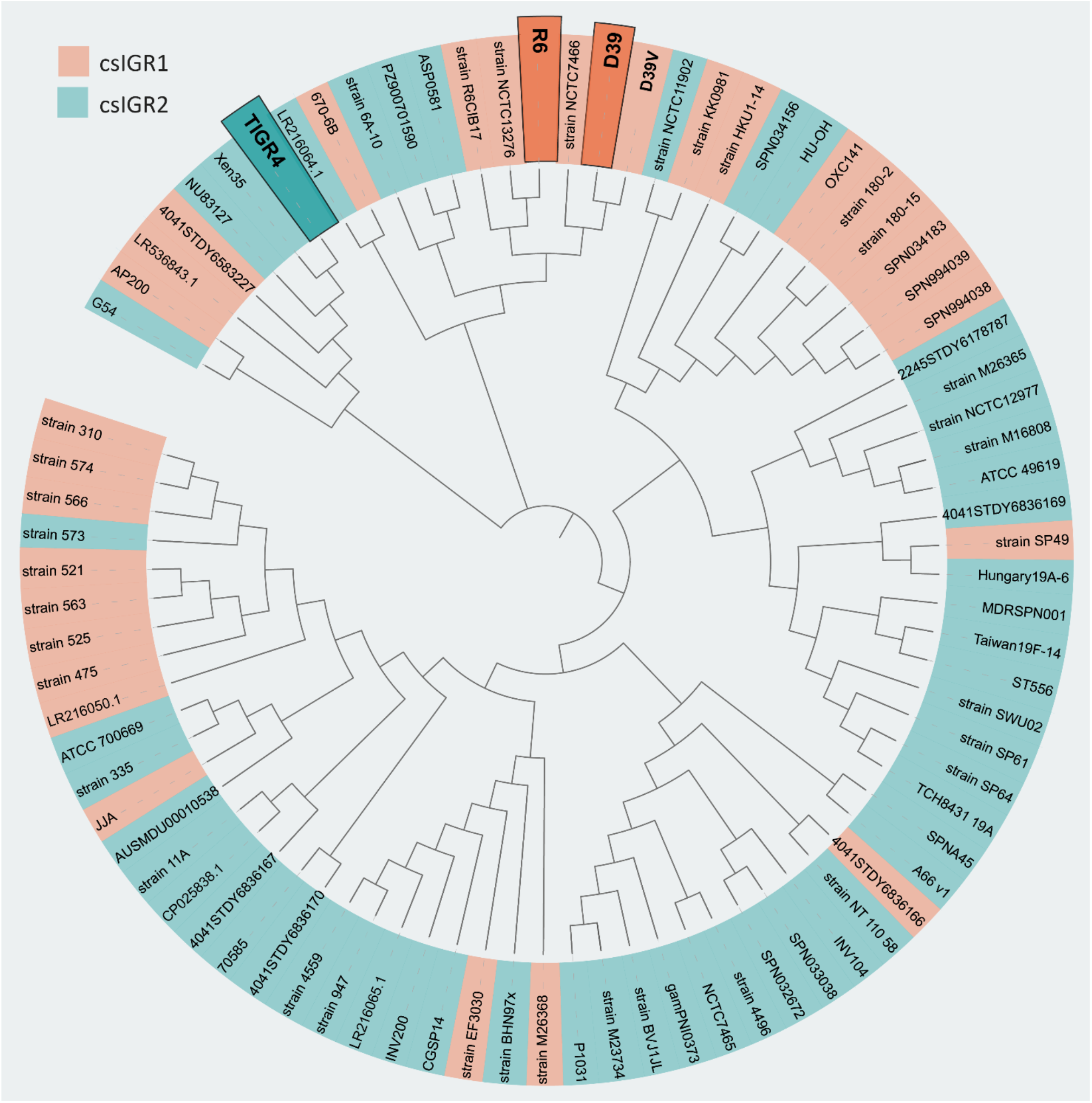
Unrooted phylogenetic tree of the 84 S. pneumoniae strains used in this study. The tree is based on SNPs in the core genes. The shading of each label indicates the presence of csIGR1 (orange) or csIGR2 (green). The tree was created using Roary, Fasttree and iTol. The S. pneumoniae strain names are specified if applicable, if no clear strain name was given the sequence ID was used.

## Discussion

Here we present the first complete pangenome of pneumococcus, spanning both genes and the non-coding IGRs. A small but conserved IGR core genome in pneumococcus was identified. We find that the pneumococcal core genome consists of 1550 unique genes and 669 IGRs, whereas the accessory genome consists of 3132 unique genes and 2683 IGRs. The number of unique genes surpasses that of unique IGR in both cases, this is unsurprising as most intraoperonic regions in pneumococcus are less than 30bp in length and are therefore disregarded in the analysis. This also means that most IGRs identified are associated with the flanking genes of operons *i*.*e*., the regulatory regions.

IGRs between two divergently transcribed genes are termed double regulatory (DR). These regions constituted a greater relative part of the core genome than the accessory genome. It is likely because meaningful regulation of two genes is harder to achieve than regulation of single genes, raising the selection threshold for the emergence of beneficial divergence. This increased selection pressure has previously been observed as purifying selection has been shown to be more prominent in DR regions than the other types [17].

Our analysis reveals that IGRs are highly conserved in pneumococcus. On average, 66% of IGRs in any isolate is shared with all other isolates and 79% of genes in any isolate is shared with all other isolates. A similar trend is seen in *S. aureus*, however, the overall number of unique IGRs is lower in pneumococcus relative to genome size. Instead, our analysis reveals that pneumococcus has several duplicates of some core IGRs across the genome, with core IGRs on average being present 1.25 times in each isolate. This trend is not seen with its core genes and is not observed in neither the core genes nor core IGRs of *S. aureus*. This suggests that pneumococcus is more rigid with its transcriptional profile as the same regulatory regions might be repeated to a greater degree than observed in *S. aureus*.

We identify a clear linkage between the core IGRs and core genes in pneumococcus, on average 81% of core IGRs are directly upstream of a core gene. This indicates that the transcriptional regulation of the core genome in pneumococcus is mostly conserved across all isolates, but to a lesser degree than seen in *S. aureus*. However, this leaves 19% of core genes being associated with accessory IGRs, indicating some plasticity to the core genome that is otherwise viewed as stable. The greatest difference seen between the two species in this regard is that core IGRs are more often associated with accessory genes in pneumococcus. This might be explained by pneumococcus retaining multiple copies of some core IGRs, making them associated with both core and accessory genes, though this remains to the investigated.

Surprisingly, only 3 switches were detected in pneumococcus and only one of these were flanked by core genes. In another study, the same analysis was done on a collection of *E. coli* genomes and 61 switches were detected [22]. This indicates that regulatory switching does not play a major role in pneumococcal disease. It is possible that regulatory switching is more prominent in *E. coli* as it inhabits a great number of different niches compared to pneumococcus [2-26].

Our results show that the pneumococcal core genome is less stable than previously thought. While there is indeed a stable reservoir of highly conserved core genes, their flanking IGRs, which contain most of the regulatory regions responsible for controlling the transcription of these core genes show greater plasticity. We believe that future studies will benefit from viewing the genes as a “package” with their upstream IGR, as even core genes maintain different regulatory regions within the pneumococcal species.

## Materials and methods

### Genomes

All 84 complete *S. pneumoniae* genomes available from the National Center for Biotechnology Information, GenBank resource was downloaded in raw FASTA format. Additionally, 84 randomly selected *S. aureus* genomes were retrieved for comparison with *S. pneumoniae* (12/5/2021). Genomes were then annotated with Prokka, using the standard parameters of the software [27]. The genomes used are listed in Appendix 3.

### Pangenome creation

Initially a pangenome of the coding sequences (CDS) was created using Roary [8]. Then a complementary pangenome of the IGRs was created using Piggy, an intergenic pangenome analysis tool that emulates Roary [22]. Some steps were taken to ensure comparability between the outputs of the software. Roary was set to cluster CDSs with -e -n (to perform alignments using MAFFT [28]), -i 90 (90% sequence identity cut-off) and -s (to not split paralogs into separate clusters). The settings for running Piggy were set at the standard parameters of the software, except for -len_id 10 (the minimum percentage of length identity to form a cluster). The length identity was reduced for comparability with Roary, as gene clusters generated by Roary only require a sequence length identity of 120bp for clustering CDSs, thus the len_id of 10 is recommended by the creators of Piggy for Roary consistency as IGRs are not erroneously placed into separate clusters [8-22]. The randomly assigned locus tags provided by Prokka were translated when necessary, by aligning the GFF files of the Genbank annotated files and Prokka output.

### Core gene and core IGR linkage analysis

The linkage of core genes and core IGRs was quantified using R. The gene_presence_absence file from Roary and the IGR_presence_absence file from Piggy was loaded as dataframes in R. For each gene and IGR cluster in the files their status as a core or accessory gene was identified and assigned. For each genome all IGRs were paired with their upstream gene. Thus, NR regions were removed from the dataset and both flanking genes for DR regions were analyzed separately, if any of the two genes were core, the DR IGR was assigned as flanking a core gene.

### Switched intergenic regions analysis

For identification of switched IGRs, a separate analysis using Piggy was performed with -len_id 90 (the minimum percentage of length identity to form a cluster). This was done to perform a more strict analysis of the IGRs, as the higher threshold for forming a cluster ensured that homologue IGRs were not identified as switched IGRs [22]. IGR switches were identified using the “gene-pair” method of Piggy, here two or more different IGR sequences that occupy the same space between a specific gene pair are analyzed. The candidate IGR sequences are then aligned with BLASTN with low complexity filtering turned off and if there are no significant matches between the IGR they are identified as “switched”. If there is a significant match Piggy aligns the sequences using MAFFT and provides the nucleotide identity of the alignments.

The identified switch was validated manually with a blastn against all the genomes (data not shown).

### Phylogenetic analysis

A phylogenetic tree of the strains included in this study was created, based on single nucleotide polymorphisms (SNPs) in the core genes. Roary was run separately with the same settings as previously mentioned with the exception of -e (Core gene alignment with PRANK) [8]. This produced a highly accurate alignment of the core genes within the pangenome. FastTree was then run to infer an approximately-maximum-likelihood phylogenetic tree based on SNPs within the core genes [29]. The resulting newick file was then visualized using iTol and exported to Adobe Illustrator [30].

### Pangenome wide association study

To identify whether any accessory genes were significantly associated with any of the csIGR alleles, a pangenome-wide association study was performed using Scoary [31]. A trait matrix was created as an input for Scoary, indicating which of the two csIGRs alleles were present in which genomes. Scoary then sorted the accessory genome provided by the gene_presence_absence file from Roary, scoring each accessory gene according to their co-occurrence with each csIGR.

