## Appendix 2 for "Adding context to the pneumococcal core genes – a bioinformatic analysis of the intergenic pangenome of *Streptococcus pneumoniae*"

>csIGR1

AATTTTCTTTAAAAGAGTTTCTTTTTATACTTTTCTGAAGTGGTGACGGACGTCAGCAAAGTCCTTCGGACTTTCATGAC
TAAAATTTGAGCCTAAGGTCTCAAATTTTCCGCAGTTGGTACAATCACTTGTACCAACTTACACCACAGCGAAAAGTAT
TCTCTATGGGGCTCGCCTTGCTCGCCCAAATTCAGACAACCCTTTCCTTTCTGTGC

>csIGR2

AATTTTCTTCCTTGTTTTTTGATAGTTATTTTAGGTGGTGAGGGACGGTCGGAACTAATTTTCCAAAGATTTCATCTTTG
AAAAATGGATTTCCTGACCTCACAATTGAGCCTTGGTCTCAATTGTGCCCCCTTGCCAATCTACAGATTGGCAATCACA
CCACGGCAATAACTATCGCTATGTGAGCTCACTTTGTTCGCTCAGCGATTTTGTT
