## Appendix 3 for "Adding context to the pneumococcal core genes – a bioinformatic analysis of the intergenic pangenome of *Streptococcus pneumoniae*"

genomes

| | Genbank_genome | Strain |
|---|---|---|
| 1 | GCA_000006885.1 | TIGR4 |
| 2 | GCA_000007045.1 | R6 |
| 3 | GCA_000014365.2 | D39 |
| 4 | GCA_000018965.1 | 70585 |
| 5 | GCA_000018985.1 | JJA |
| 6 | GCA_000019005.1 | P1031 |
| 7 | GCA_000019025.1 | Taiwan19F-14 |
| 8 | GCA_000019265.1 | Hungary19A-6 |
| 9 | GCA_000019825.1 | G54 |
| 10 | GCA_000019985.1 | CGSP14 |
| 11 | GCA_000026665.1 | ATCC_700669 |
| 12 | GCA_000146975.1 | AP200 |
| 13 | GCA_000147095.1 | 670-6B |
| 14 | GCA_000180515.2 | SPNA45 |
| 15 | GCA_000196595.1 | TCH8431/19A |
| 16 | GCA_000210935.1 | INV200 |
| 17 | GCA_000210955.1 | OXC141 |
| 18 | GCA_000210975.1 | INV104 |
| 19 | GCA_000210995.1 | SPN034156 |
| 20 | GCA_000211015.1 | SPN034183 |
| 21 | GCA_000211035.2 | SPN994038 |
| 22 | GCA_000211055.2 | SPN994039 |
| 23 | GCA_000211075.1 | SPN032672 |
| 24 | GCA_000211095.1 | SPN033038 |
| 25 | GCA_000251085.2 | ST556 |
| 26 | GCA_000299015.1 | gamPNI0373 |
| 27 | GCA_000817005.1 | strain_NT_110_58 |
| 28 | GCA_001255215.1 | A66_v1 |
| 29 | GCA_001457635.1 | NCTC7465 |
| 30 | GCA_001896045.1 | strain_SP49 |
| 31 | GCA_001896065.1 | strain_SP61 |
| 32 | GCA_001896085.1 | strain_SP64 |

| | | |
|---|---|---|
| 33 | GCA_001902455.1 | strain_SWU02 |
| 34 | GCA_002357995.1 | strain_KK0981 |
| 35 | GCA_002813535.1 | strain_BHN97x |
| 36 | GCA_002813955.1 | strain_11A |
| 37 | GCA_002843545.1 | Xen35 |
| 38 | GCA_002947575.1 | strain_335 |
| 39 | GCA_003003495.1 | strain_D39V |
| 40 | GCA_003351525.1 | strain_M23734 |
| 41 | GCA_003351645.1 | strain_M16808 |
| 42 | GCA_003351705.1 | strain_M26365 |
| 43 | GCA_003351725.1 | strain_M26368 |
| 44 | GCA_003354825.1 | strain_SPN_XDR_SMC1710-32 |
| 45 | GCA_003609935.1 | MDRSPN001 |
| 46 | GCA_003966485.1 | ATCC_49619 |
| 47 | GCA_003966505.1 | HU-OH |
| 48 | GCA_003966525.1 | NU83127 |
| 49 | GCA_003967155.2 | ASP0581 |
| 50 | GCA_003994915.1 | strain_HKU1-14 |
| 51 | GCA_004291255.1 | strain_EF3030 |
| 52 | GCA_004331935.1 | strain_521 |
| 53 | GCA_008253725.1 | strain_R6CIB17 |
| 54 | GCA_009664475.1 | strain_AUSMDU00010538 |
| 55 | GCA_011694695.1 | strain_PZ900701590 |
| 56 | GCA_013047165.1 | strain_6A-10 |
| 57 | GCA_015838875.1 | strain_310 |
| 58 | GCA_015839035.1 | strain_475 |
| 59 | GCA_015839235.1 | strain_521 |
| 60 | GCA_015839415.1 | strain_525 |
| 61 | GCA_015839575.1 | strain_566 |
| 62 | GCA_015839775.1 | strain_573 |
| 63 | GCA_015839915.1 | strain_574 |
| 64 | GCA_015840115.1 | strain_563 |
| 65 | GCA_017569245.1 | strain_BVJ1JL |
| 66 | GCA_900475305.1 | strain_NCTC7466 |

| | | |
|---|---|---|
| **67** | GCA_900475515.1 | strain_NCTC13276 |
| **68** | GCA_900475805.1 | strain_NCTC11902 |
| **69** | GCA_900476445.1 | strain_4041STDY6836167 |
| **70** | GCA_900476455.1 | strain_4041STDY6836169 |
| **71** | GCA_900476465.1 | strain_4041STDY6836170 |
| **72** | GCA_900476475.1 | strain_4041STDY6583227 |
| **73** | GCA_900476505.1 | strain_4041STDY6836166 |
| **74** | GCA_900618545.1 | strain_4496 |
| **75** | GCA_900618555.1 | strain_947 |
| **76** | GCA_900618575.1 | strain_180-2 |
| **77** | GCA_900618585.1 | strain_180-15 |
| **78** | GCA_900622505.2 | strain_4559 |
| **79** | GCA_900636585.1 | strain_NCTC12977 |
| **80** | GCA_900692935.1 | isolate_GPS_US_PATH396-sc-2[illegible] |
| **81** | GCA_900693025.1 | strain_2245STDY6178787 |
| **82** | GCA_900693055.1 | isolate_55896440-41bd-11e5-9[illegible] |
| **83** | GCA_900693075.1 | isolate_569492b0-41bd-11e5-9[illegible] |
| **84** | GCA_900795205.1 | isolate_b04a6400-1f66-11e7-b9 |
